# Hypothalamic neurosecretory protein GM causes fat deposition and suppresses gonadal maturation in Japanese quail

**DOI:** 10.64898/2026.08.24.746428

**Authors:** Masaki Kato, Eiko Iwakoshi-Ukena, Megumi Furumitsu, Yuki Narimatsu, Chisato Yatsuda, Yoshiaki Nakamura, Kazuyoshi Ukena

## Abstract

**Introduction:** Central regulation of energy homeostasis is essential for balancing lipid storage and reproductive investment; however, the hypothalamic factors governing this trade-off remain incompletely defined in birds. Neurosecretory protein GM (NPGM), an 83–amino acid hypothalamic factor, was investigated for its role in energy allocation during sexual maturation in Japanese quail (*Coturnix japonica*).

**Methods:** Male and female quails at the onset of sexual maturation received chronic intracerebroventricular administration of NPGM for 13 days via osmotic pumps, during which their body mass, food intake, and water intake were monitored daily. At the endpoint, peripheral tissue and muscle masses, serum metabolite levels (glucose, fatty acids, triglycerides, testosterone, and 17β-estradiol), hepatic triglyceride content, and gene expression profiles of hypothalamic feeding/reproductive genes and hepatic/adipose lipid metabolic genes were evaluated.

**Results:** NPGM increased subcutaneous and abdominal fat in both sexes and was associated with suppressed gonadal maturation, as indicated by reduced testicular mass relative to body mass and lower testosterone levels in males, as well as a trend toward reduced ovarian mass and lower 17β-estradiol levels in females. Sex-dependent metabolic phenotypes emerged: males exhibited increased body mass gain, hyperphagia, elevated water intake, enlarged liver, pancreas, and heart, higher serum and hepatic triglyceride levels, increased hepatic *SCD1* expression, and reduced hepatic *CGI-58*, *PPARγ*, *SLC2A2*, and *CD36*. In contrast, females showed fat accumulation without hyperphagia or hepatic triglyceride elevation, accompanied by reduced hepatic *VTG2* and *APOV1* and decreased adipose *ATGL*, *LPL*, and *FATP*. Hypothalamic *AGRP* expression decreased in males, whereas both *NPY* and *AGRP* decreased in females.

**Discussion:** These findings demonstrate that central NPGM shifts energy allocation from reproduction toward lipid storage through sex-dependent endocrine and metabolic mechanisms, identifying NPGM as a neuroendocrine regulator of energy allocation during sexual maturation in Japanese quails.

## 1 Introduction

Energy homeostasis is tightly regulated by the central nervous system (CNS) through the integration of nutritional, hormonal, and environmental signals (Kleinridders et al., 2009; Timper and Brüning, 2017). In vertebrates, hypothalamic neuropeptides play essential roles in controlling feeding behavior, energy expenditure, and lipid metabolism, thereby coordinating energy allocation among competing physiological processes (Meister, 2007; Gan et al., 2024). Notably, in animal species that exhibit reproductive behaviors or migration in response to seasonal changes, such as variations in photoperiod or temperature, the influence of CNS energy allocation is particularly evident (Shinomiya et al., 2014). Among these physiological processes, reproduction is one of the most energetically demanding events in the life cycle. The onset of sexual maturation is accompanied by substantial changes in body composition and metabolic status, requiring a coordinated redistribution of energy resources from somatic growth and storage toward gonadal development and reproductive activities (Veldhuis et al., 2005). Therefore, precise central regulation of energy allocation is essential for the successful progression of reproductive maturation. Despite the importance of energy allocation during sexual maturation, the neuroendocrine mechanisms that govern whether energy is allocated to lipid storage or reproductive development remain poorly understood. For example, alterations in 17β-estradiol levels across the reproductive cycle have been linked to cyclic changes in food intake and body fat accumulation in both mammals and birds, suggesting a tight interaction between reproductive endocrine signals and energy homeostasis (Mauvais-Jarvis et al., 2013). Therefore, identifying hypothalamic factors involved in energy allocation is crucial for understanding how animals balance energy reserves and expenditure for reproduction.

The hypothalamus is a major feeding center containing nuclei that express orexigenic and anorexigenic factors (Meister, 2007; Roger et al., 2022). Disruption of feeding regulation in the hypothalamus can lead to overeating or anorexia, resulting in obesity or emaciation (Timper and Brüning, 2017; Gan et al., 2024). While many feeding-related factors are conserved between mammals and birds, some neuropeptides and hormones are known to have opposite effects, and the complete mechanisms of energy metabolism in the avian hypothalamus remain unclear (Tachibana and Tsutsui, 2016). In recent research, we identified a novel cDNA encoding the precursor of a small secretory protein found in the chick hypothalamus (Shikano et al., 2018). This protein, consisting of 83 amino acids, has been designated as neurosecretory protein GM (NPGM) (Shikano et al., 2018). The *NPGM* gene is well conserved across vertebrates, including chickens, quails, rats, and humans (Shikano et al., 2018; Kato et al., 2024; He et al., 2026). Our previous investigations used *in situ* hybridization and immunostaining to identify the localization of NPGM in chicken and quail brains. In chickens, NPGM-expressing cells are situated in the infundibular nucleus (IN) and the medial mammillary nucleus (MM) of the hypothalamus (Shikano et al., 2018). In Japanese quail, NPGM-expressing cells are localized in the MM, and neural fibers project to the median eminence (ME). Previous studies have reported the physiological functions of NPGM (Kato et al., 2024). In birds, chronic intracerebroventricular (i.c.v.) administration of NPGM induced fat accumulation without overeating in neonatal chicks (Kato et al., 2021). In mammals, acute i.c.v. injection of NPGM or stimulation of NPGM-expressing neurons promoted feeding behavior in mice (Martinez et al., 2023; Narimatsu et al., 2023). As indicated by previous studies, NPGM is considered a regulatory factor in energy storage. Furthermore, chicken NPGM expression is upregulated in response to various stressors, including fasting, inflammation, and social isolation, suggesting a role for NPGM in regulating energy metabolism under stress (Kato et al., 2022).

*NPGM* has a paralogous gene, *NPGL,* encoding neurosecretory protein GL (NPGL), which was identified prior to *NPGM* (Ukena et al., 2014). In chicks, NPGM- and NPGL-expressing neurons partially co-localize within the hypothalamic infundibular nucleus (IN), whereas in mice, NPGL is expressed in the arcuate nucleus (Arc) and projects to anorexigenic proopiomelanocortin (POMC) neurons (Matsuura et al., 2017; Shikano et al., 2018). Furthermore, chronic i.c.v. administration of NPGL and *Npgl* gene overexpression in the hypothalamus induced obesity in chicks and rodents, whereas *Npgl*/*Npgm* double knockout mice exhibit a lean phenotype characterized by reduced food intake and increased energy expenditure (Iwakoshi-Ukena et al., 2017; Shikano et al., 2018, 2019, 2026; Narimatsu et al., 2025, 2026). Collectively, these findings suggest that NPGM and NPGL constitute an independent peptide family involved in the central regulation of energy metabolism.

However, it remains unclear how NPGM administration affects energy allocation during sexual maturation, when energy distribution undergoes significant changes. Japanese quail (*Coturnix japonica*), like chickens, are domesticated birds, but because they can undergo rapid sexual maturation through simple photoperiod manipulation and retain seasonal migratory behavior, they are an appropriate model organism for this study (Marasco et al., 2021; Sur et al., 2025). In this study, chronic i.c.v. administration of NPGM was performed in both male and female Japanese quail, and its effects on phenotype, as well as on the expression of genes related to feeding, reproduction, and lipid metabolism, were analyzed.

## 2 Materials and methods

### 2.1 Animals

Fertilized Japanese quail (*Coturnix japonica*) eggs were sourced from the Japanese Avian Bioresource Project Research Center at Hiroshima University and incubated in a laboratory incubator. Once hatched, quails were kept in a windowless environment maintained at 25°C, with a 16-hour light period followed by 8 hours of darkness, and were provided with ad libitum access to water and commercial feed (Dream Co., Ltd., Aichi, Japan). At 3 weeks of age, the sex of quails was identified by genotyping, after which they were placed in individual cages. The experimental protocols adhered to the Guide for the Care and Use of Laboratory Animals issued by Hiroshima University (Higashi-Hiroshima, Japan), and were approved by the university’s Institutional Animal Care and Use Committee (permit number C22-27).

### 2.2 Production of NPGM

Quail NPGM was produced through fluorenylmethyloxycarbonyl (Fmoc) chemistry using a peptide synthesizer (Initiator+ Alstra™; Biotage, Uppsala, Sweden), in accordance with our earlier procedure (Masuda et al., 2015; Kato et al., 2021). The amino acid sequence of NPGM was derived from our previous research (Kato et al., 2024). High-performance liquid chromatography (HPLC) verified that the synthesized NPGM had a purity exceeding 95%. The lyophilized NPGM was weighed with an analytical balance (AP125WD; Shimadzu, Kyoto, Japan).

### 2.3 Chronic i.c.v. administration of NPGM

The NPGM dosage was set at 15 nmol/day. This dosage was chosen based on previous studies involving NPGM and galanin-like peptide (Rich et al., 2007; Iwakoshi-Ukena et al., 2017; Kato et al., 2021) and was maintained to ensure consistency with earlier experiments on juvenile chicks (Kato et al., 2021). NPGM was prepared in a solution of 30% propylene glycol. Male and female quails, five weeks old and at the onset of puberty, received i.c.v. administrations of either the vehicle (30% propylene glycol) or 15 nmol/day NPGM through an administration cannula (model 328OP; Plastics One, Roanoke, VA) linked to an Alzet mini-osmotic pump (model 2002, delivery rate 0.5 μl/h; DURECT Corporation, Cupertino, CA). The cannula tip was positioned in the lateral ventricle at the following coordinates: 2.0 mm anterior to the lambda, 1.0 mm lateral to the midline, and 5.5 mm beneath the skull surface. The osmotic mini-pumps were placed subcutaneously in the neck region. Daily measurements of body mass, food intake, and water intake were taken between 8:00 and 10:00 throughout the study. After 13 days, quails were euthanized by decapitation between 12:00 and 15:00. The masses of subcutaneous fat, abdominal fat, gizzard fat, reproductive organs, liver, pancreas, heart, pectoralis major and minor muscle, and biceps femoris muscle were recorded. The hypothalamic infundibulum, abdominal fat, and liver were promptly snap-frozen in liquid nitrogen and stored at −80°C for real-time qPCR analysis. Blood samples were collected at the end of the experiment and centrifuged at 3000×g for 15 minutes at 4°C to obtain serum.

### 2.4 Serum biochemical analysis

Serum glucose levels were measured using the GLUCOCARD G+ meter (Arkray, Kyoto, Japan). Serum testosterone and 17β-estradiol were quantified using commercially available enzyme-linked immunosorbent assay (ELISA) kits (Cayman Chemical, Inc., Ann Arbor, MI). The concentrations of free fatty acids were determined using the non-esterified fatty acid (NEFA) C-Test Wako (Wako Pure Chemical Industries, Osaka, Japan). Serum triglyceride (TG) levels were assessed with the Triglyceride E-Test Wako (Wako Pure Chemical Industries).

### 2.5 Determination of triglyceride concentration in the liver

Lipid extraction from the liver was performed using a chloroform:methanol mixture (2:1) according to a previously described method (Folch et al., 1957; Narimatsu et al., 2022). TG concentrations in the liver were determined using a colorimetric assay with the Triglyceride E-Test Wako (Wako Pure Chemical Industries).

### 2.6 Real-time qPCR

RNA was extracted from the hypothalamic infundibulum using the RNAqueous Micro Kit (Thermo Fisher Scientific, Waltham, MA), from the liver using TRIzol reagent (Life Technologies, Carlsbad, CA) and Sepasol (Nacalai Tesque, Inc., Kyoto, Japan), and from abdominal fat using QIAzol lysis reagent (QIAGEN) and Sepasol (Nacalai Tesque, Inc.) following the manufacturer’s instructions. First-strand cDNA was synthesized from total RNA (1 μg) using the PrimeScript RT reagent Kit with gDNA Eraser (TaKaRa Bio, Inc., Shiga, Japan). PCR amplification was performed using THUNDERBIRD SYBR qPCR Mix (TOYOBO, Osaka, Japan) under the following conditions: 95°C for 20 s, followed by 40 cycles of 95°C for 3 s and 60°C for 30 s using a real-time thermal cycler (CFX Connect; Bio-Rad, Hercules, CA). In the liver and abdominal fat, we analyzed the genes encoding acetyl-CoA carboxylase (ACC), fatty acid synthase (FAS), stearoyl-CoA desaturase 1 (SCD1), and peroxisome proliferator-activated receptor γ (PPARγ) as lipogenic factors, along with carnitine palmitoyltransferase 1A (CPT1A), adipose triglyceride lipase (ATGL), comparative gene identification-58 (CGI-58), and peroxisome proliferator-activated receptor α (PPARα) as lipolytic factors. Insulin-like growth factor 1 (IGF-1) is included due to its role in lipolysis in the liver and its association with adipocyte proliferation or differentiation. Tumor necrosis factor-α (TNF-α) is a pro-inflammatory cytokine and is commonly used as a marker of chronic obesity-associated inflammation. Lipoprotein lipase (LPL) facilitates fatty acid uptake and triglyceride storage in adipose tissue. Solute carrier family 2 member 2 (SLC2A2) is the gene encoding glucose transporter 2, which is involved in the regulation of blood glucose levels. CD36 functions as a fatty acid transporter in the liver and adipocytes. Fatty acid transport protein (FATP) functions as a fatty acid transporter in adipocytes. Vitellogenin 2 (VTG2) and apovitellenin-1 (APOV1) are genes involved in the synthesis of yolk protein precursors in the liver in response to 17β-estradiol. In the hypothalamus, the genes encoding NPGL, NPGM, proopiomelanocortin (POMC), corticotropin-releasing factor (CRF), neuropeptide Y (NPY), agouti-related peptide (AgRP), and galanin (GAL) were selected as feeding-related factors. Gonadotropin-releasing hormone (GnRH) and gonadotropin-inhibitory hormone (GnIH) were selected as reproduction-related factors. The primer sets are listed in Table 1.

**Table 1.**
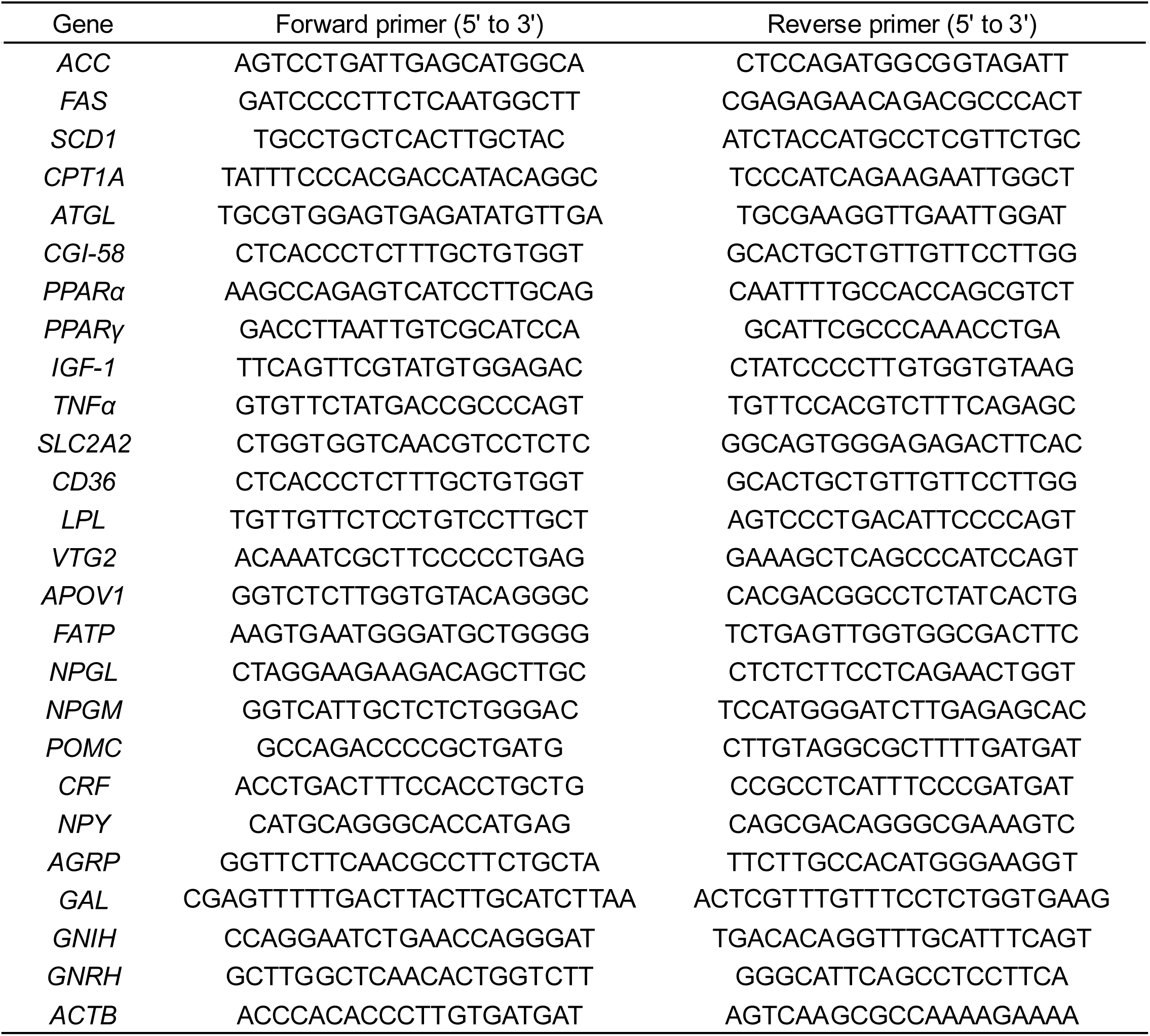
Sequences of oligonucleotide primers for real-time qPCR.

### 2.7 Statistical analysis

Differences between the two groups at the endpoint were assessed using Student’s t-test. Body mass gain and cumulative food and water intake were analyzed using two-way repeated-measures ANOVA, followed by Bonferroni’s test. The statistical significance level was set at *P* < 0.05. All data are presented as the mean ± SEM.

## 3 Results

### 3.1 Effects of chronic i.c.v. administration of NPGM in male quails

To investigate the effect of chronic i.c.v. administration of NPGM on energy metabolism in male quails, we measured body mass, food intake, and water intake for 13 days. The results showed that administration of NPGM significantly increased body mass gain, cumulative food intake, and cumulative water intake (Fig. 1A−C). Subcutaneous, abdominal, and gizzard fat mass increased in the NPGM-treated group (Fig. 1D). Among the other organs, the masses of the liver, pancreas, and heart were significantly increased (Fig. 1E). However, muscle tissue masses were unchanged (Fig. 1F). Representative photographs of abdominal fat, testes, and liver from the individuals with body masses closest to the mean in each group are shown (Fig. 1G). When testicular mass relative to body mass was compared, a significant decrease was observed in the NPGM-treated group (Fig. 1H). Serum biochemical analysis showed that the NPGM-treated group had lower testosterone levels and higher TG levels (Fig. 1I, L). In contrast, there were no changes in blood glucose or NEFA levels (Fig. 1J, K). Hepatic TG content was higher in the NPGM-treated group (Fig. 1M).

**Figure 1.**
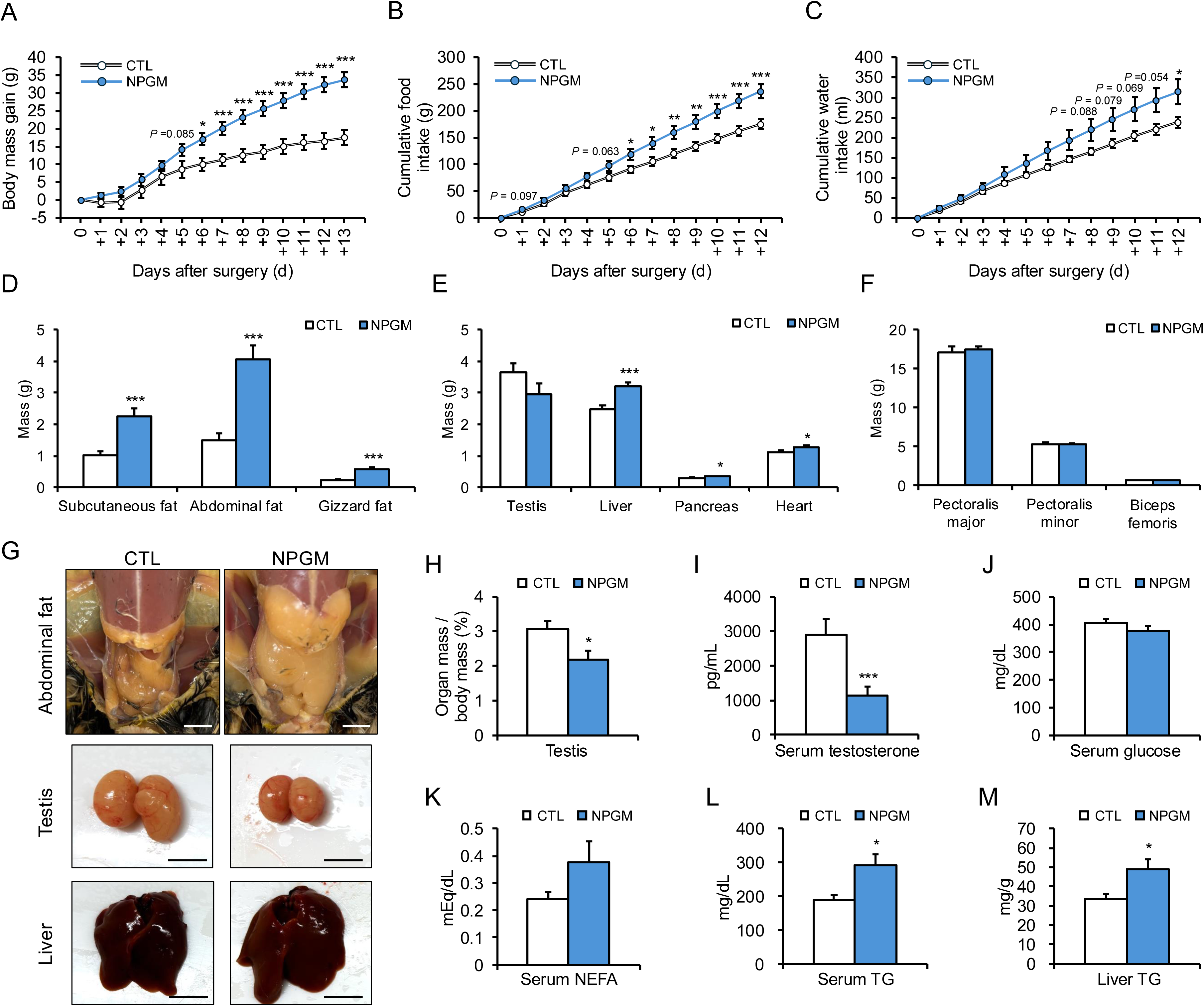
Effect of chronic i.c.v. administration of NPGM on body mass gain, food intake, and water intake in males. The results were obtained by the administration of the vehicle (control; CTL) and NPGM. (A) Body mass gain after surgery, (B) cumulative food intake, and (C) cumulative water intake. (D) Subcutaneous fat, abdominal fat, and gizzard fat mass. (E) Masses of the testis, liver, pancreas, and heart. (F) Masses of the pectoralis major, pectoralis minor, and biceps femoris. (G) Photographs of the abdominal fat, testes, and liver of individuals with body masses close to the mean for each group (Scale bar = 10 mm). (H) Testis mass per body mass. (I) Concentration of testosterone in serum. (J) Concentration of glucose in serum. (K) Concentration of non-esterified fatty acid (NEFA) in serum. (L) Concentration of triglyceride (TG) in serum. (M) Triglyceride content of the liver. Data are expressed as the mean ± SEM (n = 8). Data were analyzed using Student’s *t*-test and two-way repeated-measures analysis of variance (ANOVA). An asterisk indicates a statistically significant difference (\**P* < 0.05, \*\*\**P* < 0.005).

### 3.2 Effects of chronic i.c.v. administration of NPGM in male quails on mRNA expression of lipid metabolism-related genes

In male quails, chronic i.c.v. administration of NPGM increased liver and adipose tissue mass. Based on these results, we analyzed the mRNA expression of lipid metabolism-related genes by real-time qPCR in the liver and abdominal fat. In the liver, we assessed the expression levels of *ACC*, *FAS*, *SCD1*, *CPT1A*, *ATGL*, *CGI-58*, *PPARα*, *PPARγ*, *IGF-1*, *TNFα*, *SLC2A2*, *CD36*, *VTG2*, and *APOV1* (Fig. 2A). The NPGM-treated group showed a notable increase in *SCD1* mRNA expression, while *CGI-58*, *PPARγ*, *SLC2A2*, and *CD36* expression levels were significantly decreased (Fig. 2A). In the abdominal fat, we assessed the expression levels of *ACC*, *FAS*, *SCD1*, *CPT1A*, *ATGL*, *CGI-58*, *PPARα*, *PPARγ*, *IGF-1*, *TNFα*, *LPL*, *CD36*, and *FATP* (Fig. 2B). In the NPGM-treated group, *CGI-58* expression showed a decreasing trend, and *LPL* expression was significantly decreased (Fig. 2B).

**Figure 2.**
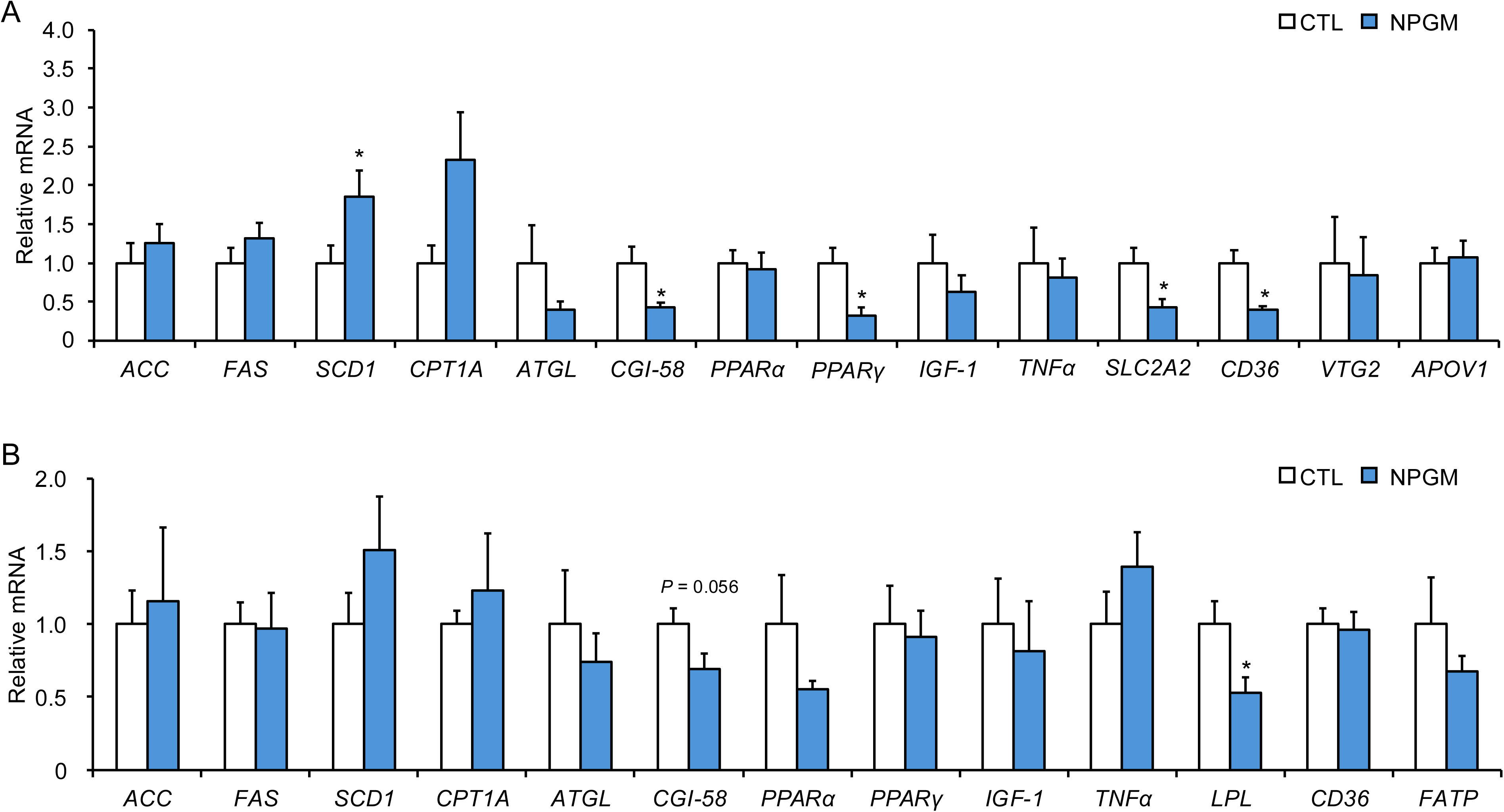
Effect of chronic i.c.v. administration of NPGM in male quails. (A) Relative mRNA expression in the liver: acetyl-CoA carboxylase (ACC), fatty acid synthase (FAS), stearoyl-CoA desaturase 1 (SCD1), carnitine palmitoyltransferase 1A (CPT1A), adipose triglyceride lipase (ATGL), comparative gene identification-58 (CGI-58), peroxisome proliferator-activated receptor α (PPARα), peroxisome proliferator-activated receptor γ (PPARγ), insulin-like growth factor-1 (IGF-1), tumor necrosis factor α (TNFα), solute carrier family 2 member 2 (SLC2A2), CD36, vitellogenin 2 (VTG2), and apovitellenin-1 (APOV1). (B) The mRNA expression in the abdominal fat: ACC, FAS, SCD1, CPT1A, ATGL, CGI-58, PPARα, PPARγ, TNFα, lipoprotein lipase (LPL), CD36, and fatty acid transport protein (FATP). Data are expressed as the mean ± SEM (n = 5-8). Data were analyzed by Student’s t-test. An asterisk indicates a statistically significant difference (\**P* < 0.05).

### 3.3 Effects of chronic i.c.v. administration of NPGM in female quails

To investigate the effect of chronic i.c.v. administration of NPGM in female quails, we measured body mass, food intake, and water intake for 13 days. The results showed that administration of NPGM did not change body mass gain, cumulative food intake, and cumulative water intake (Fig. 3A−C). However, subcutaneous and abdominal fat were significantly increased in the NPGM-treated group (Fig. 3D). Gizzard fat mass showed an increasing trend (Fig. 3D). In terms of organ mass, the mass of the ovary showed a decreasing trend, while the mass of the heart showed an increasing trend (Fig. 3E). The masses of the liver and pancreas remained unchanged (Fig. 3E). Similarly, there were no alterations in the muscle tissue masses (Fig. 3F). Representative photographs of abdominal fat, ovary, and liver from the individuals with body masses closest to the mean in each group are shown (Fig. 3G). The ratio of ovarian mass to body mass also showed a decreasing trend (Fig. 3H). Serum biochemical analysis showed that the NPGM-treated group had lower 17β-estradiol levels (Fig. 3I). In contrast, there were no changes in blood glucose, NEFA, or TG levels (Fig. 3J−L). Hepatic TG content was unchanged in the NPGM-treated group (Fig. 3M).

**Figure 3.**
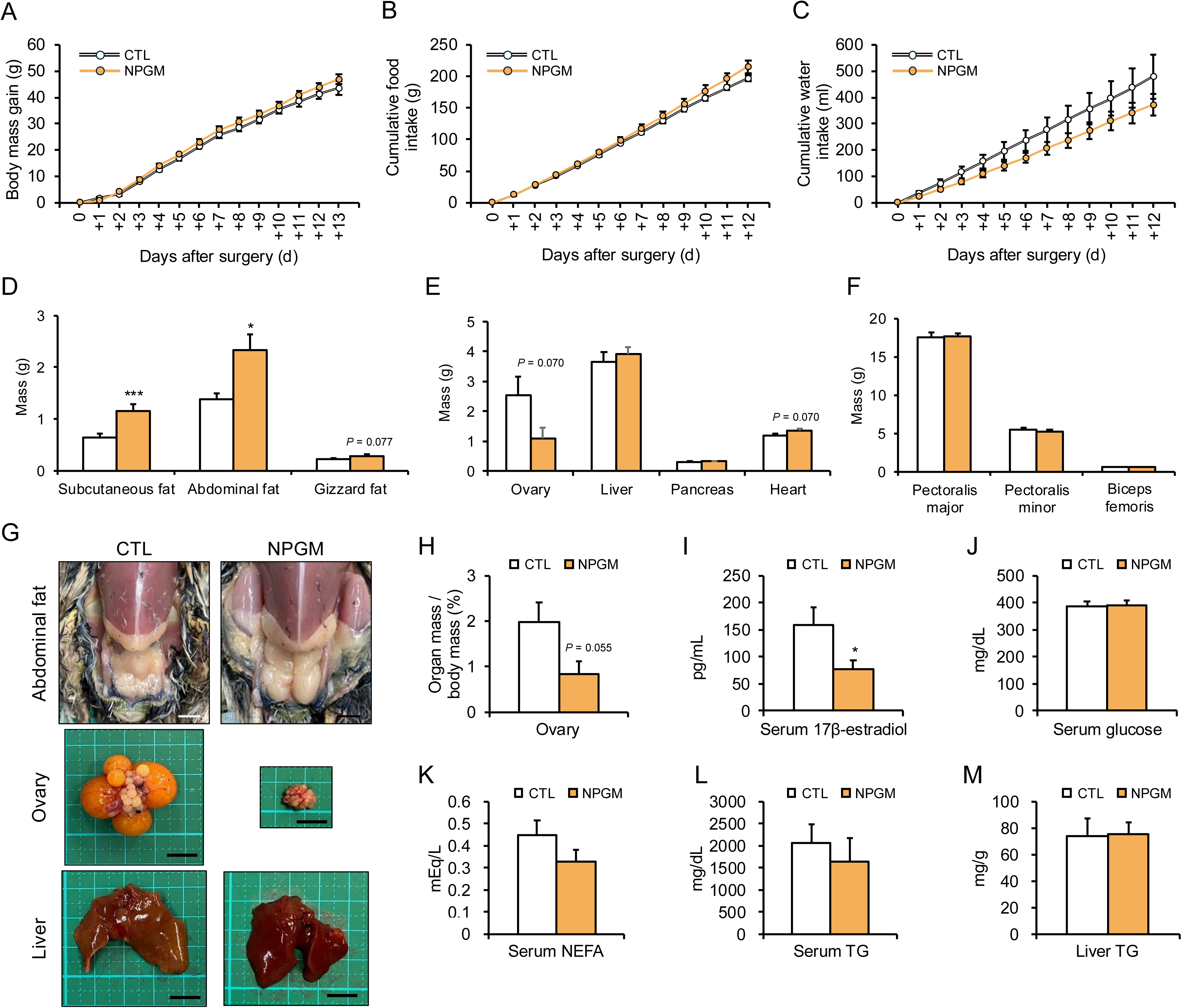
Effect of chronic i.c.v. administration of NPGM on body mass gain, food intake, and water intake in females. The results were obtained by the administration of the vehicle (control; CTL) and NPGM. (A) Body mass gain after surgery, (B) cumulative food intake, and (C) cumulative water intake. (D) Subcutaneous fat, abdominal fat, and gizzard fat mass. (E) Masses of the ovary, liver, pancreas, and heart. (F) Masses of the pectoralis major, pectoralis minor, and biceps femoris. (G) Photographs of the abdominal fat, ovary, and liver of individuals with body masses close to the mean for each group (Scale bar = 10 mm). (H) Ovarian mass per body mass. (I) Concentration of 17β-estradiol in serum. (J) Concentration of glucose in serum. (K) Concentration of NEFA in serum. (L) Concentration of TG in serum. (M) Triglyceride content of the liver. Data are expressed as the mean ± SEM (n = 10). Data were analyzed using Student’s *t*-test and two-way repeated-measures analysis of variance (ANOVA). An asterisk indicates a statistically significant difference (\**P* < 0.05, \*\*\**P* < 0.005).

### 3.4 Effects of chronic i.c.v. administration of NPGM in female quails on mRNA expression of lipid metabolism-related genes

In female quails, chronic i.c.v. administration of NPGM increased adipose tissue mass. Based on this result, we analyzed the mRNA expression of lipid metabolism-related genes by real-time qPCR in the liver and abdominal fat. In the liver, *VTG2* and *APOV1* mRNA expression were significantly decreased in the NPGM-treated group (Fig. 4A). In the abdominal fat, *ATGL*, *LPL*, and *FATP* were significantly decreased (Fig. 4B).

**Figure 4.**
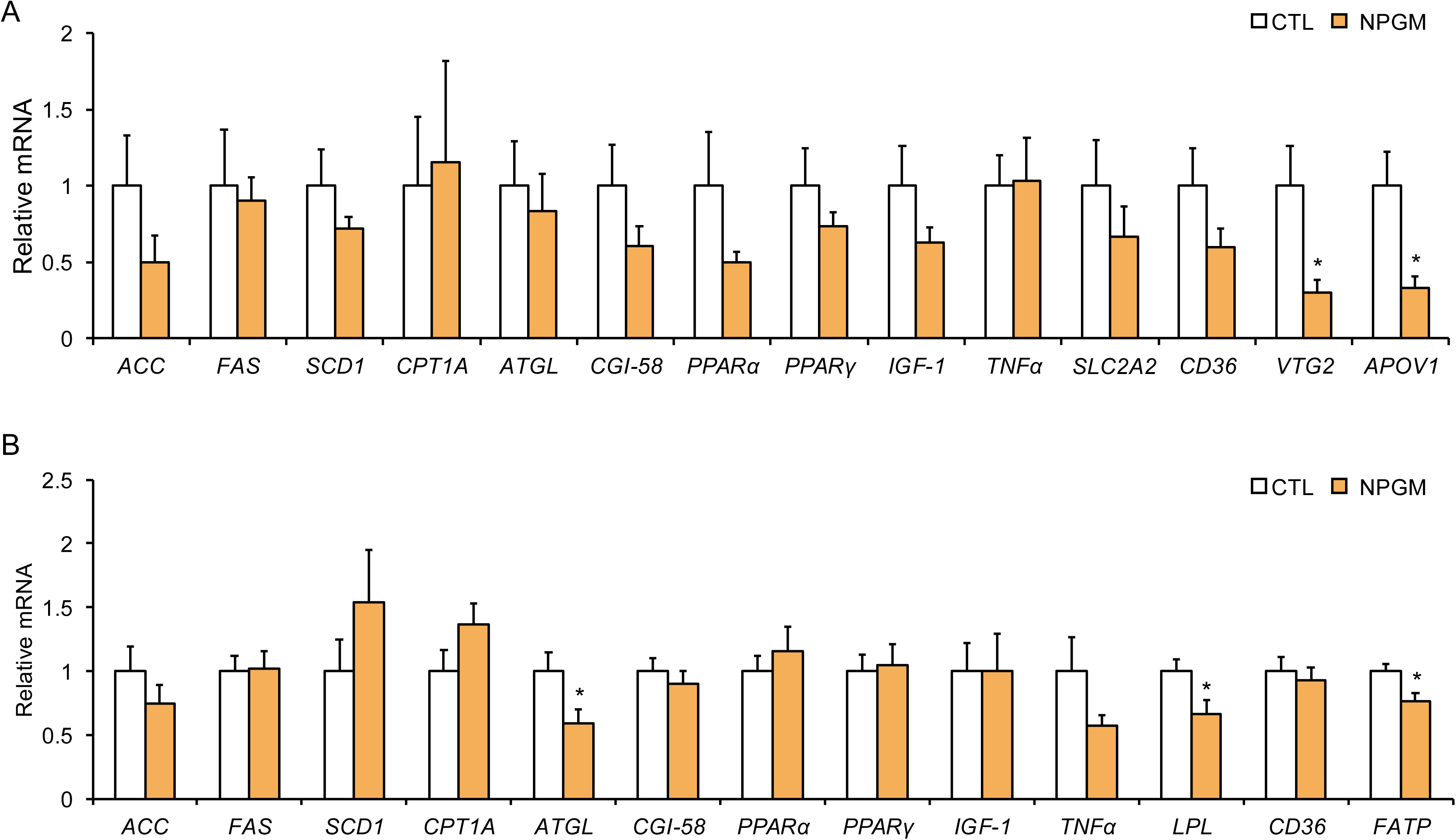
Effect of chronic i.c.v. administration of NPGM in female quails. (A) Relative mRNA expression in the liver: ACC, FAS, SCD1, CPT1A, ATGL, CGI-58, PPARα, PPARγ, IGF-1, TNFα, SLC2A2, CD36, VTG2, and APOV1. (B) The mRNA expression in the abdominal fat: ACC, FAS, SCD1, CPT1A, ATGL, CGI-58, PPARα, PPARγ, TNFα, LPL, CD36, and FATP. Data are expressed as the mean ± SEM (n = 10). Data were analyzed by Student’s t-test. An asterisk indicates a statistically significant difference (\**P* < 0.05).

### 3.5 Effects of chronic i.c.v. administration of NPGM on mRNA expression of feeding-related factors and reproduction-related hormones in the hypothalamus

As fat accumulation and delayed maturation of the gonads were observed in both male and female quail, the gene expression levels of feeding-related factors (*NPGL*, *NPGM*, *POMC*, *CRF*, *NPY*, *AGRP*, *GAL*) and reproduction-related hormones (*GNIH*, *GNRH*) expressed in the hypothalamus were analyzed by qPCR. The administration of NPGM led to a reduction in mRNA expression of *AGRP* in male quails (Fig. 5A), while both *NPY* and *AGRP* mRNA expression levels were decreased in female quails (Fig. 5B). The mRNA expression levels of *GNIH* and *GNRH* remained unchanged in both sexes (Fig. 5A, B).

**Figure 5.**
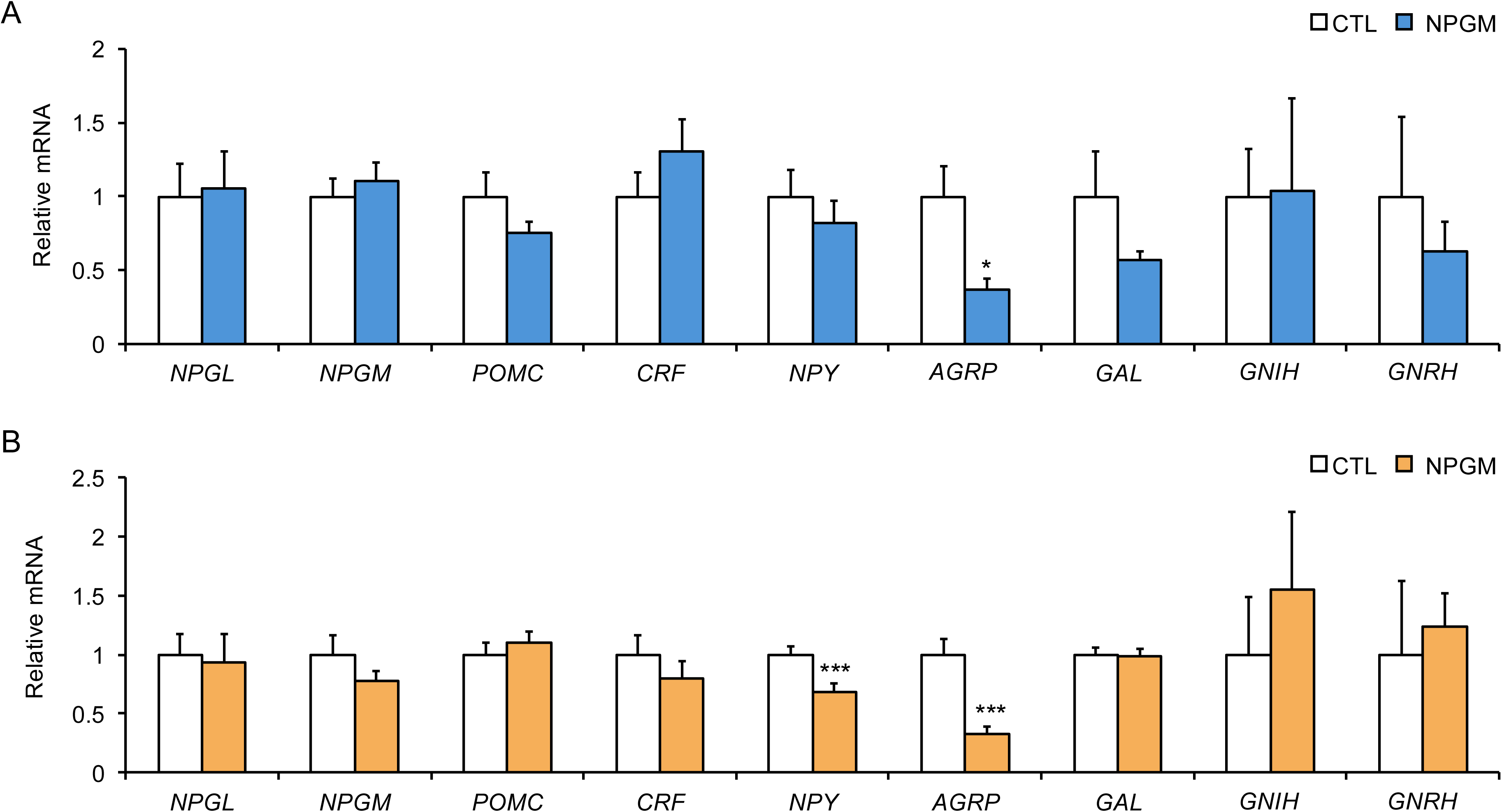
Effect of chronic i.c.v. administration of NPGM on the mRNA expression of hypothalamic factors in males (A) and females (B). Neurosecretory protein GL (NPGL), neurosecretory protein GM (NPGM), pro-opiomelanocortin (POMC), corticotropin-releasing factor (CRF), neuropeptide Y (NPY), agouti-related peptide (AGRP), and galanin (GAL), gonadotropin-inhibitory hormone (GNIH), and gonadotropin-releasing hormone (GNRH). The results were obtained by the administration of the vehicle (control; CTL) and NPGM. Data are expressed as the mean ± SEM (n = 8–10). Data were analyzed by Student’s t-test. Asterisks indicate statistically significant differences (\**P* < 0.05).

## 4 Discussion

In this study, we investigated the effects of a novel hypothalamic small protein, NPGM, on the body composition of both male and female quails by chronic i.c.v. administration for 13 days. As common phenotypes in both sexes, NPGM administration promoted fat accumulation and inhibited gonadal maturation. In contrast, feeding behavior and gene expression related to lipid metabolism differed between sexes, suggesting that the mechanism of action of NPGM varies by sex.

First, we focused on sex differences in the effects on feeding behavior. This was because NPGM is expressed in the hypothalamus, the feeding center, and is a paralog of NPGL, an orexigenic factor (Ukena et al., 2014; Shikano et al., 2018). In the case of male quails, NPGM increased body mass, food intake, and fat accumulation. The results of this study were similar to those reported in mammals such as mice. In male mice, acute i.c.v. administration of NPGM enhances feeding behavior (Martinez et al., 2023), and it has been reported that pharmacological manipulation specific to NPGM-expressing neurons leads to short-term increases in feeding behavior (Narimatsu et al., 2023). In addition, fasting experiments have been shown to increase *NPGM* mRNA expression in the hypothalamus of chicken, quail, and mice (Kato et al., 2022, 2024; He et al., 2026). These reports suggest that NPGM is an orexigenic factor in both mammals and birds. Furthermore, this hypothesis is supported by the finding that NPGM in mice is expressed in a subset of AgRP neurons and belongs to an evolutionarily older lineage than AgRP (He et al., 2026). On the other hand, in chickens, NPGM is co-expressed with the anorexigenic neurotransmitter histamine in the MM, and acute i.c.v. administration of NPGM to male neonatal chicks has been shown to suppress feeding behavior (Shikano et al., 2018). Thus, although *NPGM* gene expression shows a similar response to fasting in birds and mammals, pharmacological administration of NPGM exerts opposite effects on feeding behavior in these two groups. Whether the relationship between feeding behavior and NPGM is evolutionarily conserved warrants further careful investigation in future studies.

In this study, gene expression analysis in the hypothalamus revealed that the expression of *AGRP*, an orexigenic neuropeptide, was significantly reduced in both sexes following NPGM administration. These results suggest that the suppression of *AGRP* expression may be a common response to NPGM administration. On the other hand, regarding the effects on feeding behavior, while food intake increased in males, no significant difference in food intake was observed in females, despite the promotion of fat accumulation. These results indicate that sex-specific neural circuits and endocrine factors other than AgRP are likely involved in the sex differences in the effects of NPGM on feeding behavior. There have been reports of functional differences between males and females in hypothalamic factors regulating food intake in mammals. For example, the orexigenic actions of NPY and AgRP are known to be modulated by sex hormones, particularly estradiol, resulting in sex-dependent differences in feeding behavior and metabolic regulation in mice (Olofsson et al., 2009; Torres Irizarry et al., 2022). Likewise, anorexigenic POMC neurons exhibit enhanced activity in females compared with males in mice (Wang et al., 2018). Similarly, sex differences in the expression of feeding-related factors have been observed in chickens, with higher hypothalamic *NPY* and *AGRP* expression in males than in females, whereas no sex differences were observed in *POMC* and *CART* expression (Caughey et al., 2018). These findings suggest that hypothalamic feeding circuits are differentially regulated between sexes. In contrast, our previous study demonstrated that basal *NPGM* gene expression does not differ between male and female quails (Kato et al., 2024). Therefore, the sex-dependent effect of NPGM on food intake observed in the present study may reflect differences in the neural networks or hormonal environments through which NPGM exerts its action. Future studies are needed to characterize the neural connectivity between NPGM-expressing neurons and feeding-regulatory neuronal populations in a sex-specific manner.

In the present study, chronic i.c.v. administration of NPGM increased adipose tissue mass while decreasing gonadal mass and circulating sex hormone levels in both male and female quails. We initially hypothesized that the increased adiposity and suppressed gonadal development induced by chronic i.c.v. administration of NPGM were mediated through alterations in the hypothalamic-pituitary-gonadal (HPG) axis. Specifically, we expected that NPGM would regulate the mRNA expression of GnIH and GnRH, two key hypothalamic neuropeptides controlling reproductive function (Schwanzel-Fukuda and Pfaff, 1989; Tsutsui et al., 2000). However, gene expression analysis revealed no significant changes in the expression levels of either *GNIH* or *GNRH* in either sex. These findings suggest that the effects of NPGM on lipid metabolism and reproductive function are unlikely to be mediated through direct regulation of GnIH and/or GnRH pathway. Although the downstream signaling pathway of NPGM remains unclear, previous studies have suggested several possible mechanisms. Transcriptomic analysis of the mouse-derived hypothalamic neuronal cell line mHypoE-46 suggested that NPGM is co-expressed in a subset of AgRP neurons (He et al., 2026). In addition, GPR83 has been identified as a candidate receptor for NPGM by an in vitro binding assay using HEK293T cells (Li et al., 2023). However, direct anatomical evidence demonstrating the colocalization of NPGM with AgRP neurons or the projection relationship between NPGM neurons and GPR83-expressing neurons has not yet been established in birds.

Because our findings suggest that the effects of NPGM are unlikely to be mediated through direct regulation of GnIH and GnRH, we considered the possibility that NPGM may function not only as a regulator of lipid metabolism but also as a key factor coordinating the balance between reproduction and energy storage. Although NPGM promoted adiposity in both sexes, the underlying mechanisms may differ between males and females. In birds, the liver is the primary organ responsible for lipid synthesis and lipid transport (Hermier, 1997). During the laying period, lipids synthesized in the liver are transported to developing follicles together with yolk precursor proteins, including VTG2 and APOV1 (Li et al., 2014). In the present study, liver mass was unchanged in females; however, the expression levels of *VTG2* and *APOV1* were significantly decreased, accompanied by reductions in serum 17β-estradiol levels and ovarian mass (Fig. 6B). These findings suggest that hepatic lipid transport for yolk formation was suppressed, resulting in the redistribution of lipids that would be incorporated into the yolk toward adipose tissue (Fig. 6B). Therefore, in females, NPGM may promote adiposity by suppressing lipid allocation to reproduction (Fig. 6B). In contrast, males exhibited increased liver mass together with elevated *SCD1* expression, whereas the expression of *CGI-58*, *PPARγ*, *SLC2A2*, and *CD36* was reduced (Fig. 6A). Moreover, serum testosterone levels and testicular mass were decreased. In humans and rodents, testosterone deficiency is recognized as a risk factor for obesity and is associated with increased visceral fat accumulation and reduced fatty acid oxidation (McInnes et al., 2012; Kelly and Jones, 2013). Therefore, the reduction in testosterone induced by NPGM administration may have contributed to adiposity (Fig. 6A). Thus, NPGM-induced fat accumulation in males may involve multiple mechanisms, including alterations in lipid metabolic pathways and reduced testosterone levels (Fig. 6A). Future studies are needed to elucidate the pathways through which NPGM regulates lipid metabolism and sex hormone synthesis.

**Figure 6.**
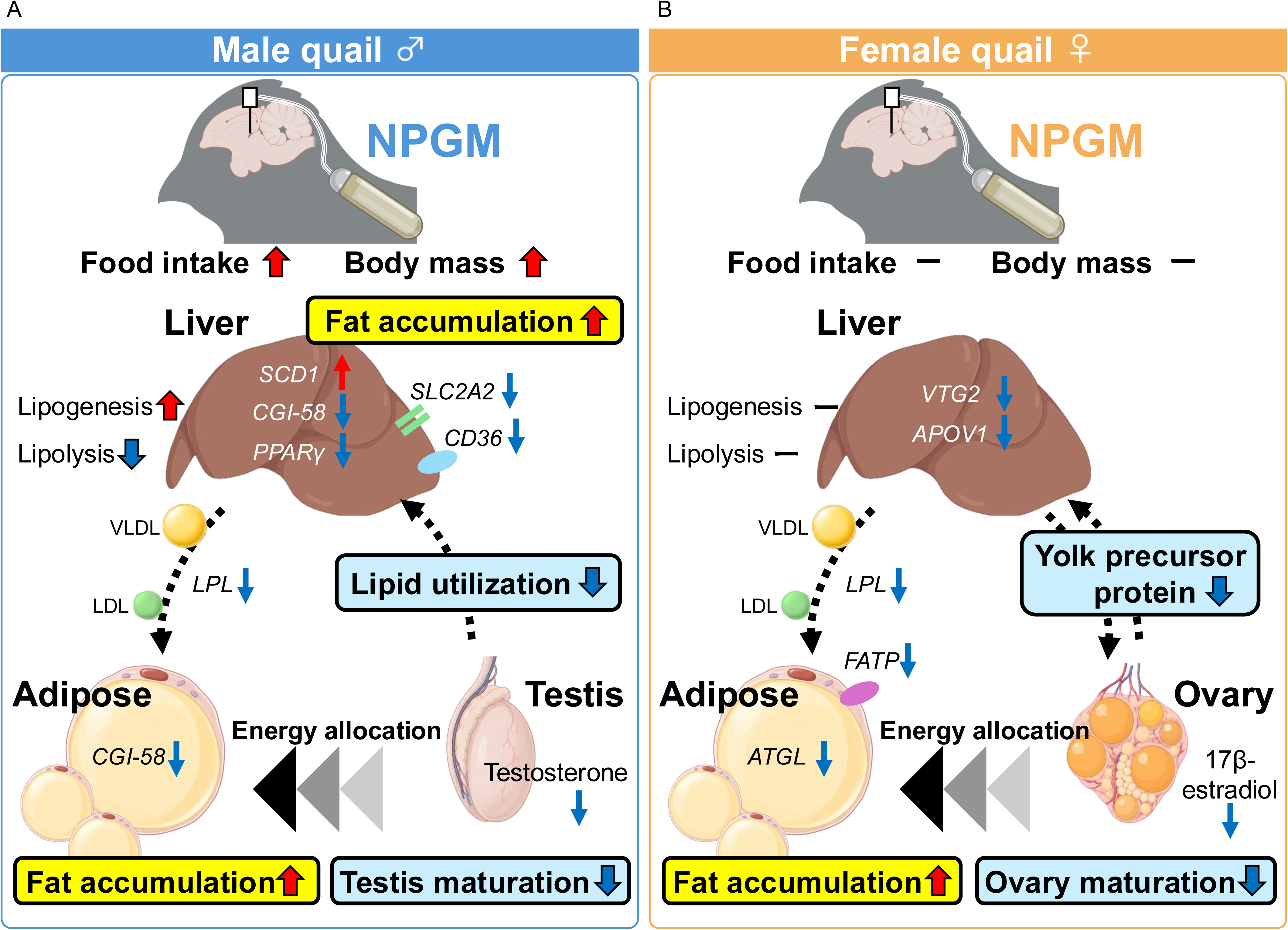
Proposed mechanisms for NPGM-induced fat accumulation and inhibition of gonadal maturation in male and female quails. (A) In male quails, NPGM induces hepatic and adipose tissue changes in lipid metabolism-related gene expression together with reduced circulating testosterone, resulting in decreased lipid utilization and increased fat accumulation. (B) In female quails, NPGM suppresses hepatic expression of the yolk precursor genes, accompanied by reduced circulating 17β-estradiol, thereby decreasing lipid allocation to developing follicles and redirecting energy toward adipose tissue. These findings support a model in which NPGM coordinates energy allocation by shifting energy allocation from reproductive investment toward lipid storage through distinct endocrine and metabolic pathways in males and females. Downward blue and upward red arrows indicate ‘downregulate’ and ‘upregulate’, respectively and other-colored arrows demonstrate downstream events. Created in BioRender. Kato, M. (2026) https://BioRender.com/cbp18w5.

Obesity in rodents and humans is associated with enhanced expression of fatty acid uptake-related factors, such as LPL and CD36, in adipose tissue, thereby promoting the uptake of lipoprotein-derived fatty acids and subsequent lipid accumulation (Picard et al., 2002; Bonen et al., 2006). LPL and CD36 are key regulators of fatty acid uptake into adipocytes in chickens (Hermier, 1997; Shu et al., 2011). However, *LPL* expression was reduced in the adipose tissue of both sexes, despite the marked increase in adipose tissue mass observed in this study. Furthermore, the expression of *FATP* in female adipose tissue was also reduced (Fig. 6B). In addition, the expression of lipolytic genes was decreased, including *CGI-58* in males and *ATGL* in females (Fig. 6A, B). These findings suggest that NPGM-induced fat accumulation is unlikely to result from enhanced fatty acid uptake, but rather from suppressed lipid mobilization and utilization (Fig. 6A, B).

Taken together, these findings support the hypothesis that NPGM functions as a regulator of energy allocation, prioritizing energy storage over reproduction through distinct molecular and endocrine mechanisms in males and females (Fig. 6A, B). In females, NPGM appears to promote fat accumulation by suppressing lipid transport to the developing yolk, whereas in males it may do so through testosterone-dependent suppression of lipid utilization (Fig. 6A, B). Consistent with this hypothesis, visceral fat mass normally decreases after the onset of egg laying in chickens as lipids are redirected toward yolk production (van Eck et al., 2023). Furthermore, in seasonally breeding species, especially mammals and birds, fat deposition is enhanced after the breeding season to prepare for migration or overwintering (Florant et al., 2004; Guglielmo et al., 2005; Sur et al., 2025). Collectively, these observations suggest that NPGM functions as a physiological regulator that temporarily suppresses reproductive investment and redirects energy toward lipid storage in preparation for future reproductive events or environmental challenges.

In conclusion, the present study demonstrates for the first time that chronic i.c.v. administration of NPGM promotes fat accumulation and inhibits gonadal maturation in quails during the process of sexual maturation (Fig. 6). This study provides new insights into the neuroendocrine mechanisms that regulate energy allocation during sexual maturation in birds. However, this study has several limitations. Because the present study employed chronic administration of exogenous NPGM, the physiological roles of endogenous NPGM remain to be determined. Therefore, future studies should investigate the physiological functions of endogenous NPGM using avian adeno-associated virus (AAV)-mediated gene overexpression and genome-editing approaches to generate loss-of-function models in Japanese quails (Matsui et al., 2012; Lee et al., 2019).

Finally, because quail are long-day breeding birds that exhibit marked photoperiodic responses and undergo seasonal migration, the physiological roles of NPGM are likely to be influenced by changes in day length. Because the present study was conducted exclusively under long-day conditions, it remains unclear whether NPGM actions are conserved across different photoperiodic states. Future studies investigating the function of NPGM under varying photoperiods, including short-day conditions, will be important for determining whether NPGM contributes to seasonal adaptation by coordinating energy allocation among lipid storage, reproduction, and migration.

## 5 Data availability statement

The original contributions presented in the study are included in the article/supplementary material, further inquiries can be directed to the corresponding author.

## 6 Ethics statement

The experimental protocols were in accordance with the Guide for the Care and Use of Laboratory Animals prepared by Hiroshima University (Higashi-Hiroshima, Japan).

## 7 Author contributions

Conceptualization, M.K. and K.U.; methodology, M.K., E.I-U., M.F., and K.U.; investigation, M.K., E.I-U., M.F., Y.N., C.Y., and K.U.; writing—original draft preparation, M.K.; writing—review and editing, M.K. Y.N., and K.U.; visualization, M.K.; project administration, K.U.; funding acquisition, M.K., E.I.-U. and K.U. All authors have read and agreed to the published version of the manuscript.

## Acknowledgments

We are grateful to Shiho Tokushima, Mayo Hirano, Natsumi Takahashi, Naoki Takamatsu, Satoru Ohira, Haruto Shirahase, Saori Harada, Haruto Takeuchi, Asma Khatun, and Shogo Moriwaki (Hiroshima University) for the experimental support.

## 10 Generative AI statement

The authors have stated that generative AI was used in the preparation of this manuscript to refine, clarify, and proofread the text.

